# Exploring the Mechanism of Na^+^/K^+^‑ATPase (NKA) and 20-HETE Ligand Interactions by in-silico modeling

**DOI:** 10.64898/2026.05.12.724327

**Authors:** Dhilhani Faleel, Ramesh Arnest, Vaishnavi Aradhyula, Satkeerth Boyapalli, Steven T. Haller, David J. Kennedy

## Abstract

The Na^+^/K^+^-ATPase (NKA) regulates ion balance in the kidney and influences cellular processes like proliferation and apoptosis through its signal transduction. The endogenous ligand 20-Hydroxyeicosatetraenoic acid (20-HETE) contributes to inflammation and fibrosis in chronic kidney disease (CKD) and inhibits NKA activity in renal tubules. However, the molecular mechanism of this interaction remains unclear.

In this study, we used *in-silico* approach to investigate the potential interaction between 20-HETE and NKA. Various ligands, including known NKA ligands such as cardiotonic steroids (CTS), 20-HETE, and negative controls, were docked using rigid and Induced Fit Docking to predict the affinity of the ligands toward NKA. Binding free energy calculations with the Prime Molecular mechanics with generalized Born and surface area (Prime MM/GBSA) tools were used to confirm the involvement of key amino acids in ligand-receptor interactions.

The docking analyses revealed that 20-HETE exhibited a binding affinity comparable to negative control, with some differences between rigid and induced fit docking. Binding free energy data highlighted key amino acids in the 20-HETE and NKA interaction. Interaction fingerprint and mutations such as Ala330Gly and Val329Ala significantly reduced binding free energy, while Thr804Ala showed a notable decrease, underscoring the potential importance of these amino acids in ligand stabilization.

These findings provide computational evidence supporting potential direct interaction between 20-HETE and NKA and identify candidate residues for future experimental validation.

## 1. Introduction

The Sodium-Potassium ATPase/Na^+^/K^+^-ATPase (NKA) stands out as a highly expressed membrane protein in mammalian cells, distinguished by its pivotal role in actively pumping ions in an energy-dependent manner.[1] Its primary function revolves around maintaining the balance of ions within cells, ensuring proper ion homeostasis.[1,2] NKA is known as a P-type membrane protein and has been the focus of extensive structural investigations due to its functional significance. The NKA is assembled from three major subunits: α-subunit, the β-subunit, and the γ-subunit. The α-subunit, the largest amongst the three, exhibits a complex structure with ten transmembrane regions (referred to as M1 to M10) spanning through the cell membrane.[1,3,4] This substantial subunit boasts a molecular weight of 112 kilodaltons (KD) and comprises 1028 amino acids (AA).[4] Within the α-subunit, two critical regions play distinct roles: a sizable cytoplasmic domain where ATP and sodium ions form essential interactions, and an extracellular region that accommodates the binding of an important steroid hormone called CTS such as ouabain and potassium ions (K^+^). Previous studies have illuminated insights into the binding of the CTS, ouabain to the NKA, revealing that it occurs while the enzyme is in the E2 state.[1] This interaction effectively stabilizes a particular conformation of the NKA, subsequently inhibiting its pumping activity. Furthermore, these investigations have highlighted the remarkable stability of this conformation, making interactions between the NKA and other proteins highly improbable.[1,2,4,5]

Apart from this canonical function the pump also completes non-canonical functions.[1] Previous studies show that while NKA can act as a pump in renal tubule cells it also acts as a receptor to which CTS can act as a ligand.[6] The key to this signaling is the NKA-Src interaction; however, chronic elevation of CTS can result in continuous activation of the NKA-Src signaling complex which leads to chronic inflammation and fibrosis in vital tissues.[7]

Evidence suggests that the arachidonic acid metabolite 20-Hydroxyeicosatetraenoic Acid (20-HETE) has a natriuretic effect and is known to inhibit NKA pumping activity in renal proximal tubules and the thick ascending loop of Henle, suggesting that 20-HETE interacts with NKA.[8] However, a detailed molecular mechanism of this interaction is not well understood. Further, 20-HETE also plays a major role in inflammation associated with CKD [9–13] Therefore, this study aimed to use an *in-silico* approach to test the hypothesis that 20-HETE directly interacts with the NKA.

Protein-ligand interactions are fundamental to essential biological processes. These interactions are primarily evaluated through binding affinity assessment, typically quantified using the dissociation constant (Kd), where a lower Kd signifies a stronger binding affinity.[14] Enabling protein-ligand interactions as a cornerstone of drug discovery and design. Numerous studies have been conducted to elucidate the critical significance of these interactions, employing structural biological techniques such as X-ray crystallography and nuclear magnetic resonance spectroscopy (NMR).[15,16] Nevertheless, these methods encounter limitations, especially when applied to membrane proteins, due to the challenges associated with crystallizing membrane-bound proteins and the band broadening effects observed in NMR. [15,16]

It is worth noting that experimentally determining protein-ligand interactions, particularly for large proteins like the NKA, is not only resource-intensive but also associated with experimental complexities. Molecular docking, on the other hand, strives to replicate the interaction between a ligand and a protein. Its objective is to forecast the binding geometry and estimate the binding strength, typically measured as binding affinity. In summary, this current study aims to demonstrate that NKA may interact with 20-HETE directly using an *in-silico* modeling approach.

## 2. Results

The proinflammatory molecule 20-HETE may play a crucial role in chronic CKD by binding with NKA. To gain deeper insights into the molecular interaction mechanism between 20-HETE and NKA, we employed *in-silico* methods, including glide docking (both rigid and induced fit docking) and prime-MM/GBSA approaches. Our study investigated three well-known CTS (Ouabagenin, Telocinobufagin (TCB), and marinobufagenin (MBG)), 20-HETE, and their analogs (5,14-20-HEDGE and 5,14-20-HEDE) as well as common vehicles such as DMSO and EtOH (**Figure 1**)

**Figure 1.**
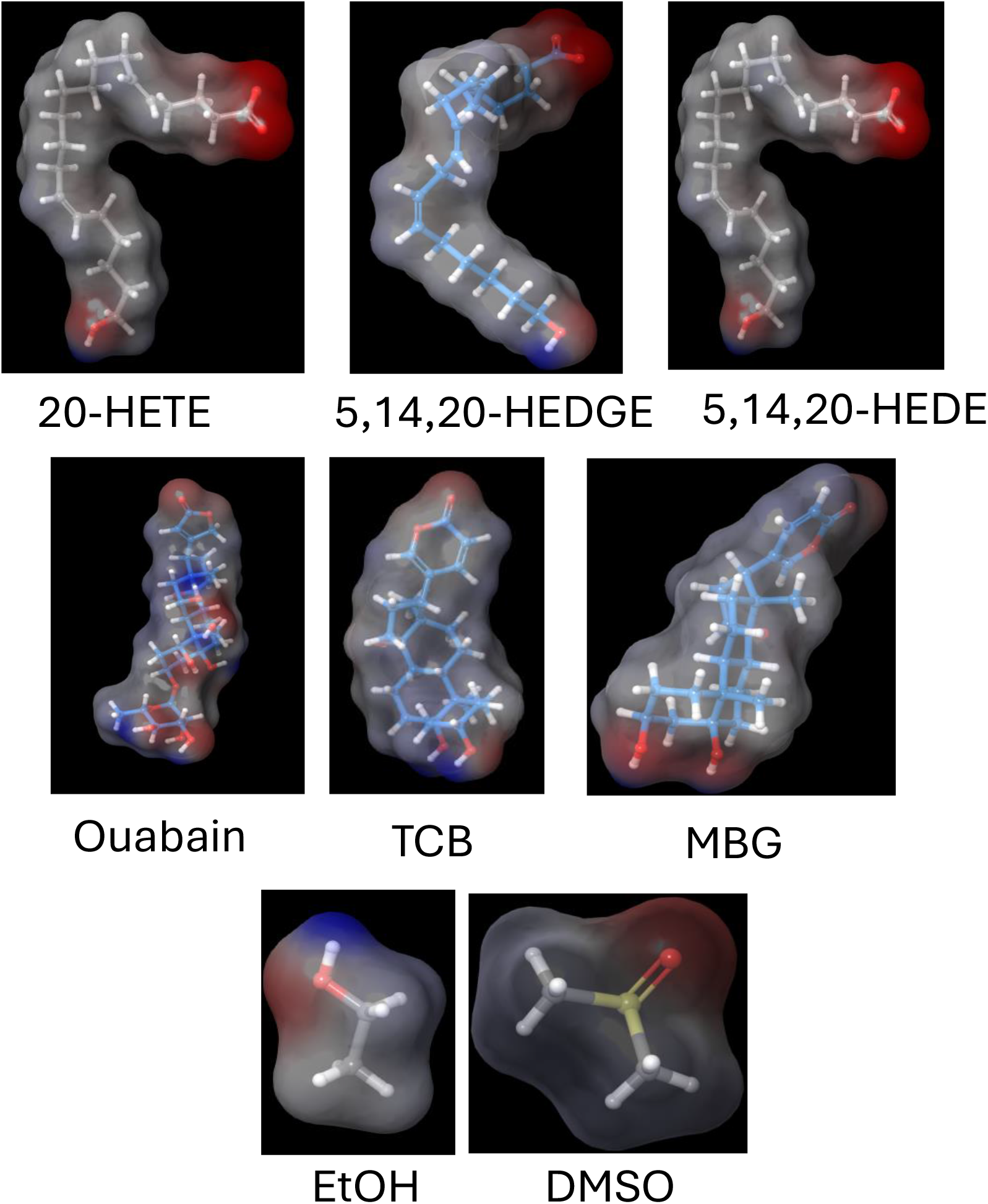
Surface representation of ligand electrostatic Potential. Using OPLS2005 Force Field (Blue: Electropositive Charge, Red: Electronegative Charge, White: Neutral

### 2.1 Docking validation by redocking

Redocking in molecular docking refers to the process of docking a ligand back into the binding site of a protein structure from which it was originally co-crystallized. Redocking serves as a validation step for the accuracy and reliability of molecular docking algorithms and scoring functions. If a docking method can successfully predict the experimental binding mode of a ligand back into its co-crystallized binding site, it provides evidence that the method is capable of reproducing known binding interactions. Further, it suggests this method could be used to establish the docking protocol of unknown or new independent datasets.[17,18]

In this study, we extracted ouabagenin from its co-crystallized structure and completed a re-docking experiment to validate localization within the NKA structure. An RMSD of 0.85 Å was obtained upon superimposed the redocked ligand with the co-crystallized ligand (**Figure 2a and 2b**). The redocked ligand shows a similar interaction profile with key binding site residues relative to the crystallographic structure (**Figure 2c and Figure S1**).[19] This shows that the prepared protein structure showed highly similar binding geometry and acceptable for further study.

**Figure 2.**
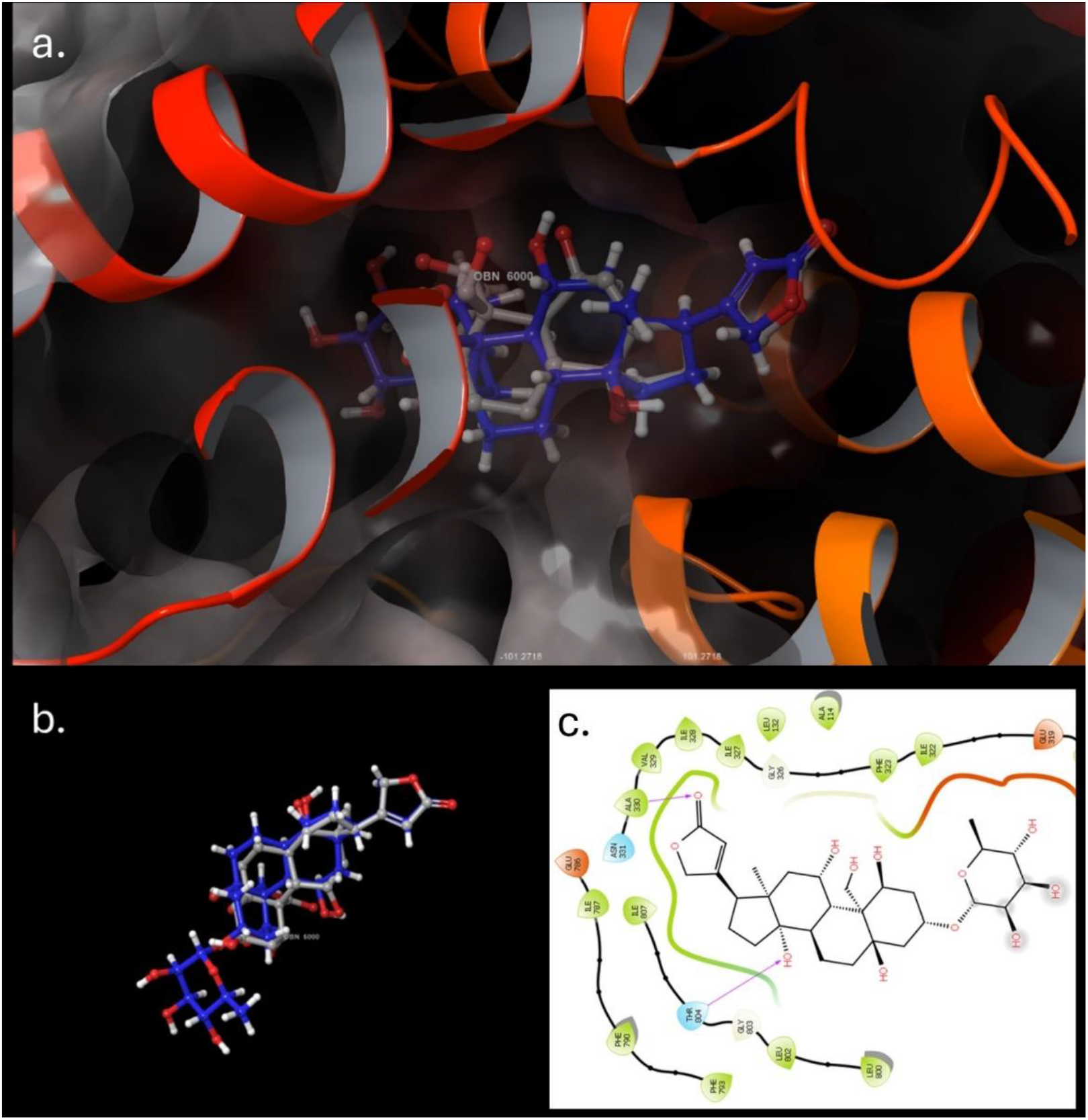
Re-docking images depicting the co-crystallized ligand in blue and the docked ligand in grey. (a) Illustrates the positioning of both the self and co-crystallized ligands within the active site. (b) Shows a superimposed 3D view of the co-crystallized and docked ligands. (c) Presents a 2D representation of the docked ouabain interaction.

### 2.2 Rigid docking scores of target molecules

A chemically divers database of urology compounds (Supplementary Table 1) was docked to Na^+^/K^+^-ATPase to establish a baseline distribution of rigid docking scores (**Figure 3**). The distribution shows a broad range of binding affinities, with most compounds clustering around −8.0 to −4.0 kcal/mol. This is consistent with high, moderate and non-specific interactions. Mapping biologically relevant ligands onto this baseline distribution revealed a clear and biologically consistent trend. CTS, including MBG (−10.2 kcal/mol), TCB (−10.1 kcal/mol), and ouabain (−10.0 kcal/mol) are in strong binding affinity region (left tail) of the distribution, establishing these metabolites as compounds with as high-affinity as Na^+^/K^+^-ATPase ligands. In contrast, negative controls such as DMSO (−2.7 kcal/mol) and EtOH (−0.8 kcal/mol) localize to the weak-binding region (right side), confirming minimal predicted interaction with the target binding pocket. Notably, 20-HETE (−5.6 kcal/mol)) and its analogs (5,14-20-HEDGE: −7.9 kcal/mol); 5,14-20-HEDE: −5.1 kcal/mol) were positioned within an intermediate region of the distribution, distinct from both CTS and negative controls. This supports 20-HETE’s interaction potential with Na^+^/K^+^-ATPase that was not likely due to random variation alone.

**Figure 3.**
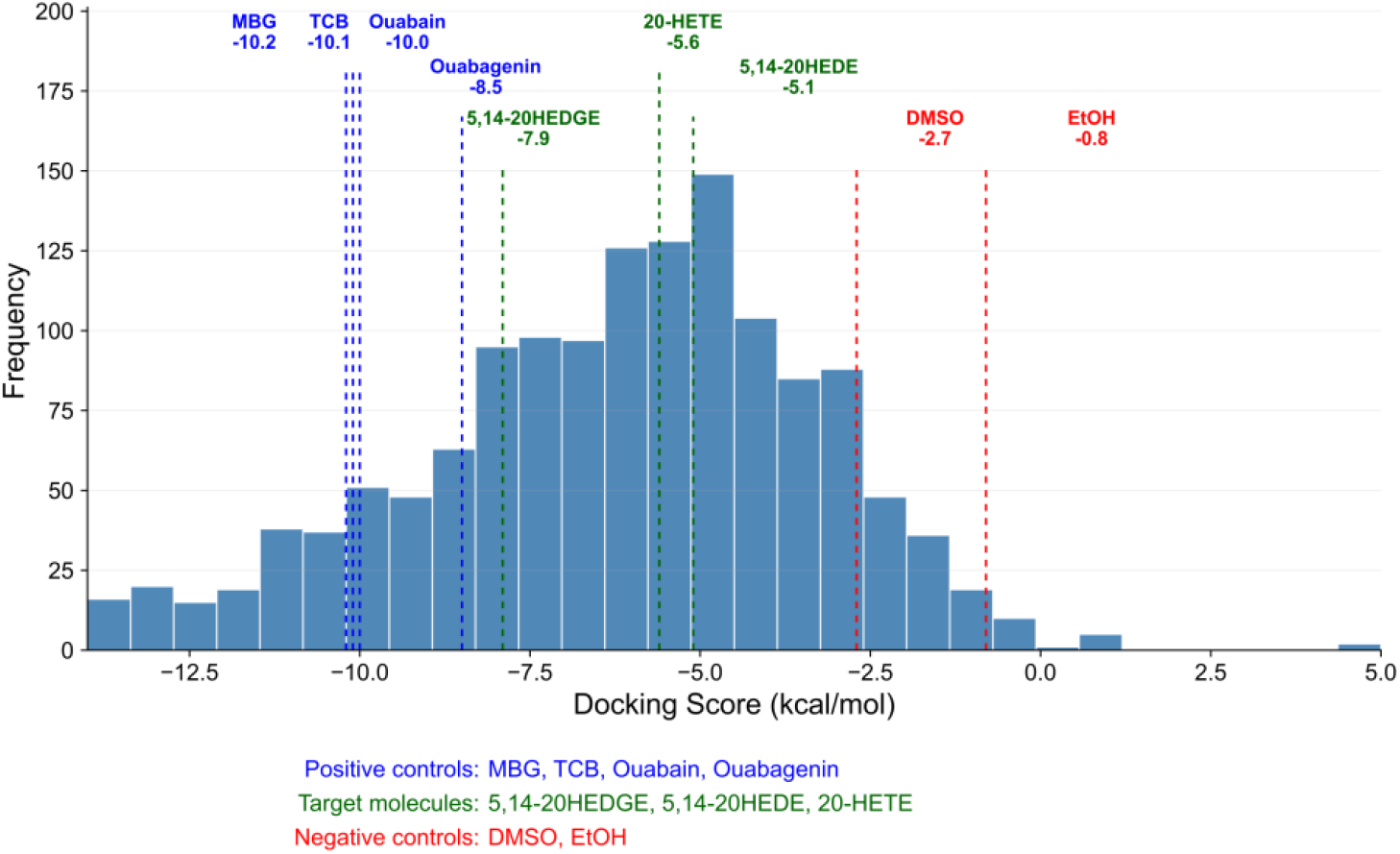
Histogram illustrates the distribution of docking scores against NKA, obtained from molecular docking of chemically divers database of urology compounds. The blue bars represent the frequency of docking scores across the background chemical space. The blue dashed lines indicate cardiotonic steroids (CTS; MBG, TCB, ouabain, and ouabagenin), the green dashed lines represent 20-HETE and its analogs (5,14-20-HEDGE and 5,14-20-HEDE), the red dashed lines indicate the negative controls (DMSO and EtOH). More negative docking scores indicate stronger predicted binding affinity toward NKA

**Figure 4.**
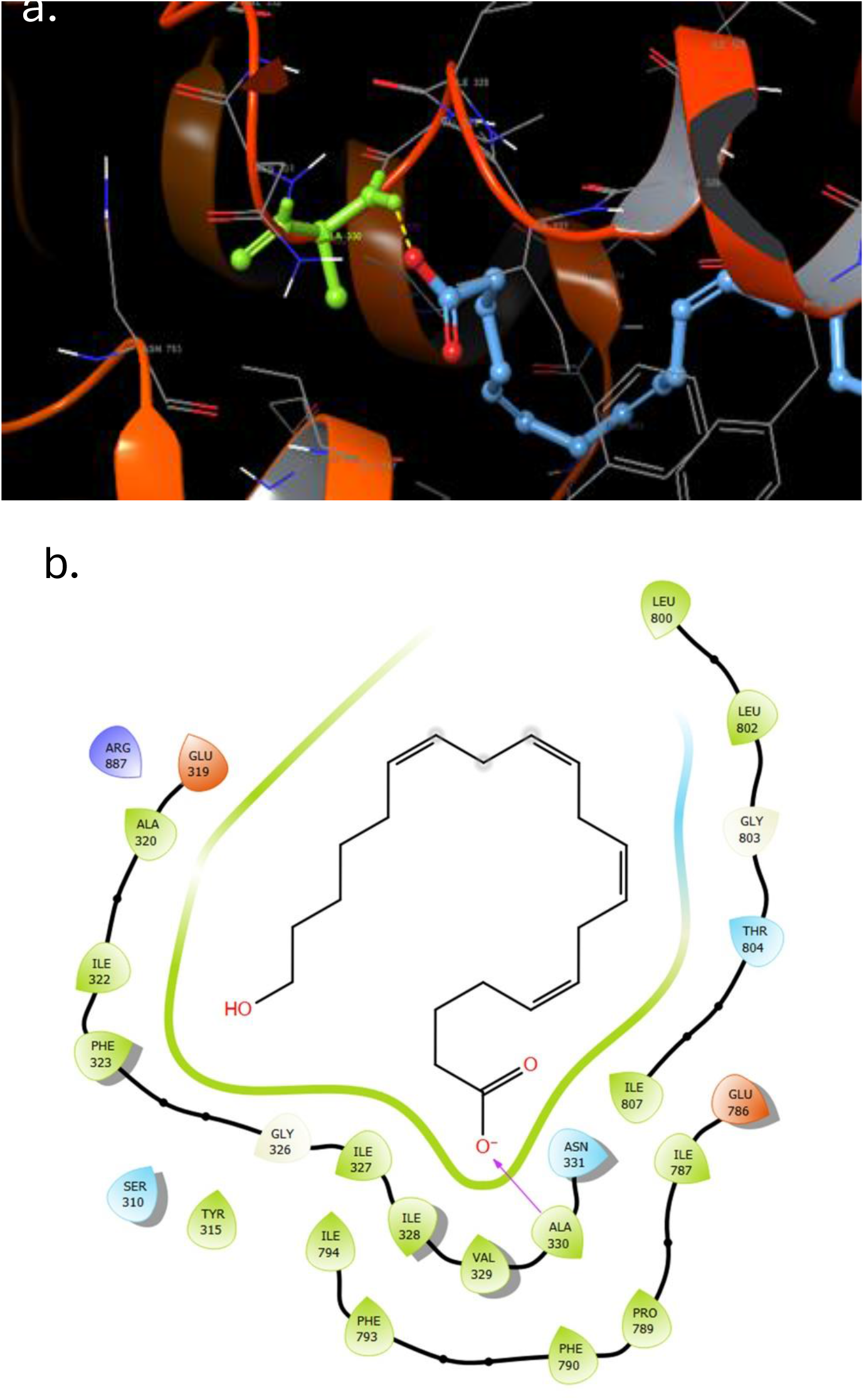
Interaction diagram between 20-HETE and NKA. a. Depicts a 3D image of the interaction, with cyan representing 20-HETE and green indicating Ala 330 of NKA. The yellow dashed line illustrates a non-classical methyl C–H···O interaction b. 2D diagram offering a more detailed view of the interaction between Ala 330 and Thr 804 of NKA with 20-HETE.

### 2.3 Induced fit docking (IFD) scores of target molecules

The Induced Fit Docking (IFD) method was employed to dynamically adjust the side chains of NKA to accommodate the ligand and simulate the protein’s conformational changes during ligand binding. This approach enhances the likelihood of successful ligand binding by allowing for molecular flexibility. IFD was conducted for all target ligands, and the resulting IFD scores were found to closely resemble the outcomes of rigid docking.

Notably, CTS exhibited the highest binding affinity in the order of Ouabain (−13.2), Ouabagenin (−10.6), TCB (−10.2), and MBG (−10), while 20-HETE and its analogs demonstrated at least two to three times higher binding affinities (5,14-20-HEDGE: −9.0, 20-HETE: −8.8, 5,14-20-HEDE: −6.5) compared to the negative controls (DMSO: −3.0, EtOH: −0.9) (**Table 1**).

**Table 1.**
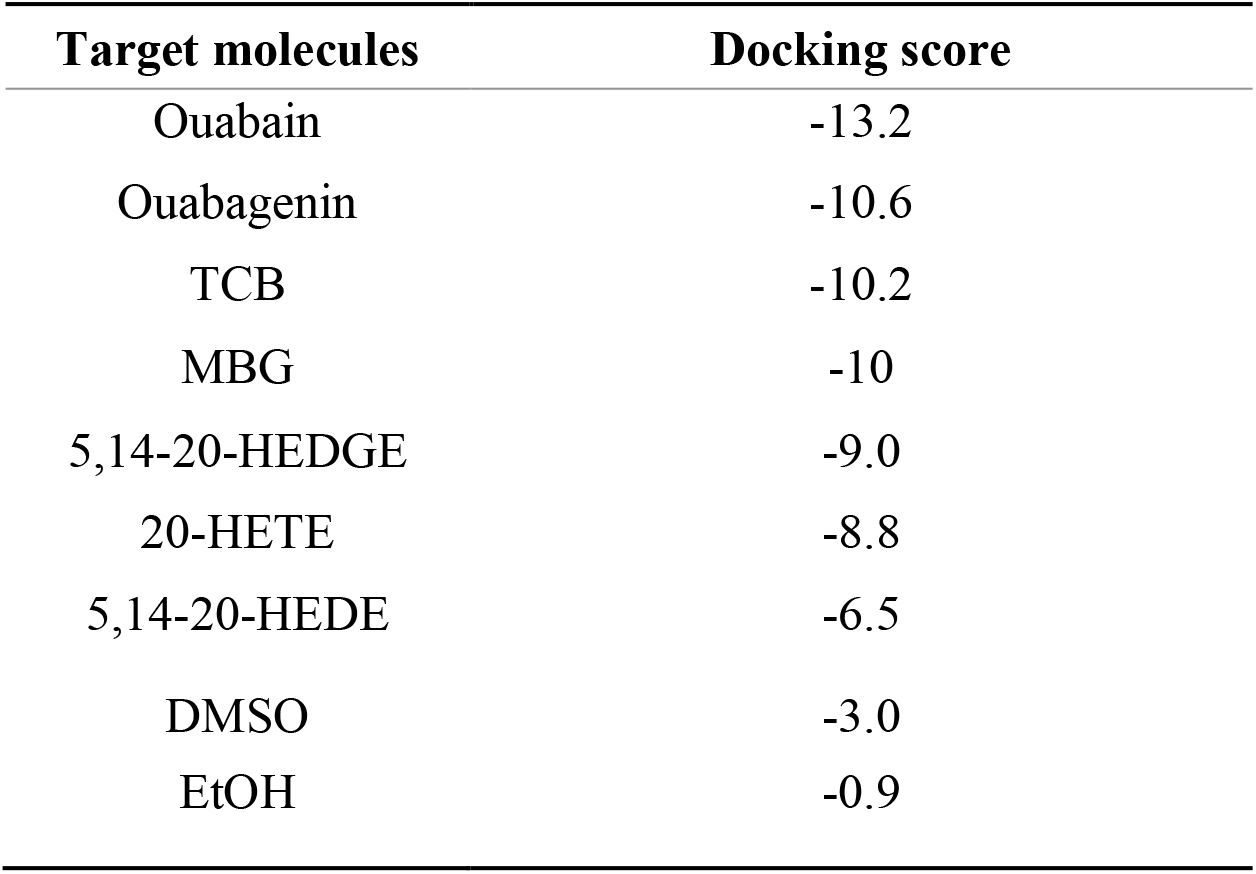
Induced fit docking scores of potential target molecules with the Na/K ATPase.

Upon closer examination, multiple significant interactions between 20-HETE and NKA were observed throughout the entire IFD simulation. These interactions predominantly comprise hydrophobic interactions. Further analysis of the interaction fingerprint data also shows several hydrophobic contacts surrounding 20-HETE within the binding pocket of NKA. Notably, the interaction of Ala330 and Thr804 with 20-HETE consistently maintained throughout the simulation, with Ala330 appears to contribute to ligand stabilization through a combination of steric positioning and a potential non-classical methyl C–H···O interaction involving its sidechain and Thr804 facilitating polar interactions[20] **(Figure 5)**

**Figure 5.**
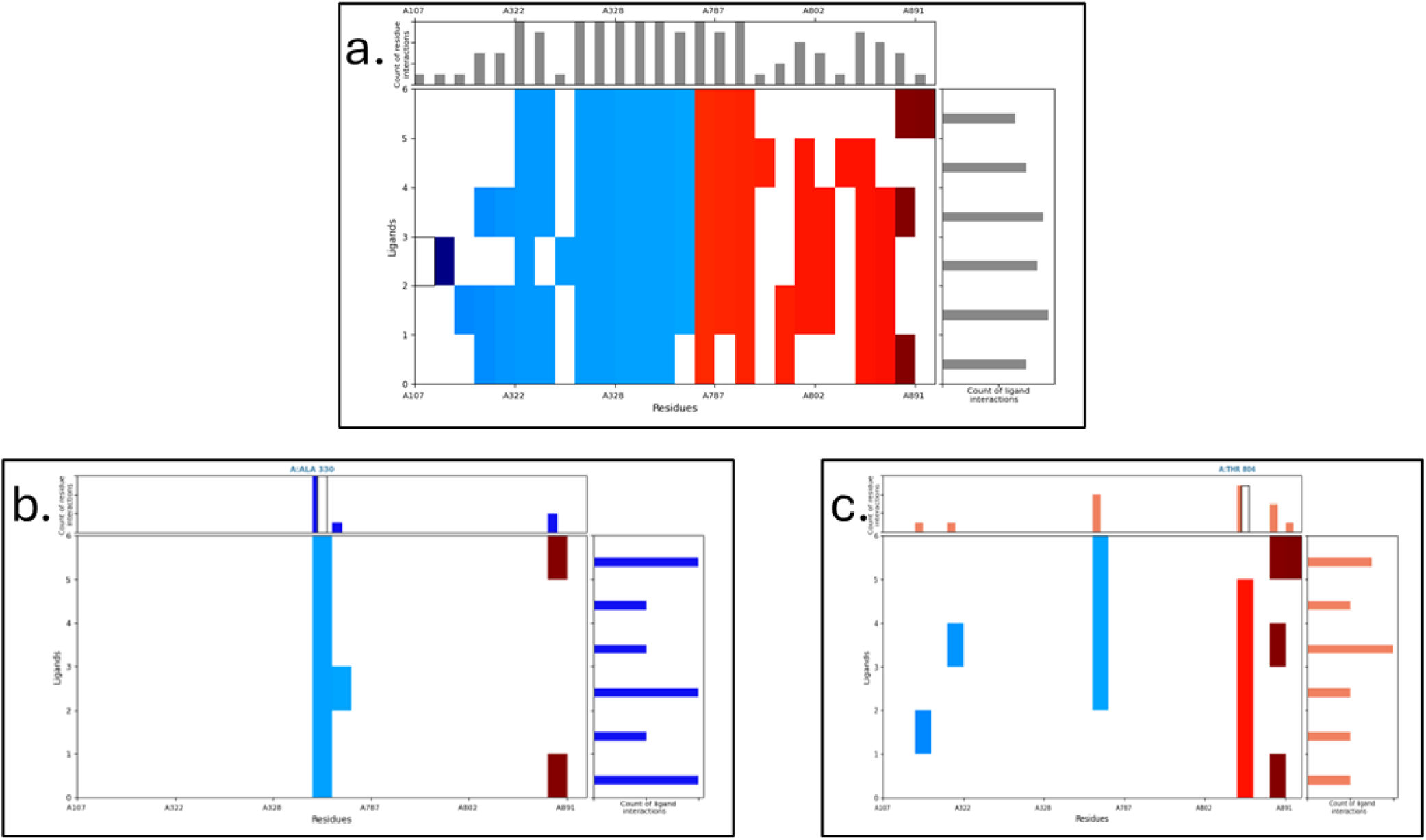
Fingerprint image depicting the IFD total simulation of 20-HETE. (a) Shows the comprehensive interaction between 20-HETE and NKA amino acids. (b) Highlights the persistent interaction of Ala 330 with 20-HETE throughout the IFD process. (c) Illustrates that Thr 804 primarily interacts with most of the poses during IFD.

### 2.4 Binding free energy calculation

Prime MM/GBSA (Molecular Mechanics/Generalized Born Surface Area) is a computational method used in molecular modeling and structural biology to estimate the binding free energy of a protein-ligand complex. Prime MM/GBSA combines molecular mechanics, which models the physical interactions between atoms, and the Generalized Born (GB) model for solvation energy to calculate the thermodynamic properties of protein-ligand interactions.[21]

In the context of molecular docking and virtual screening studies, Prime MM/GBSA is often used to refine and rank the binding affinities of ligands to a target protein. This method can provide more accurate predictions of binding energies compared to standard docking scores by considering the energetics of the solvation process and the entropic contributions to binding.[22]

The Prime MM/GBSA analysis suggest that the native NKA-ligand complex has the favorable binding free energy (−43.9556 Kcal/mol) compared to most of the mutated NKA-20-HETE complexes (Ala330=>Gly: −20.2651 Kcal/mol, Val329=>Ala: −20.6945 Kcal/mol, Glu786=>Ala: −37.817 Kcal/mol, Ile328=>Ala: −38.4154 Kcal/mol, Thr804=>Ala: −30.466 Kcal/mol, Ile327=>Ala: −42.203 Kcal/mol). However, it is interesting to note that Phe790=>ala NKA mutant has more favorable binding (46.7048 Kcal/mol) free energy than native complex **(Table 2)**. Collectively, these relatively binding free energy calculations suggest that residues Ala330, Val329, and Thr804 may contribute to stabilizing the predicted interaction of 20-HETE with NKA.

**Table 2.** Binding free energy of various mutants and native NKA in complex with 20-HETE.

| MUTATED AMINO ACIDS | ΔG BIND (KCAL/MOL) |
| --- | --- |
| Wt | -43.9556 |
| ala330=>gly | -20.2651 |
| val329=>ala | -20.6945 |
| thr804=>ala | -30.466 |
| glu786=>ala | -37.817 |
| Ile328=>ala | -38.4154 |

**Table 3.** Binding free energy Ile 327 and Phe 790 mutants and native NKA in complex with 20-HETE.

| Mutated amino acids | $\Delta G$ bind (Kcal/mol) |
| --- | --- |
| Wt | -43.9556 |
| Ile327=>Ala | -42.203 |
| Phe790=>Ala | -46.7048 |

## 3. Discussion

Chronic kidney disease (CKD) represents a major public health burden in the global population with substantially high morbidity and mortality.[23] Progression of this condition leads to end-stage kidney disease, thereby increasing the associated mortality rate. This multifactorial progressive disease is primarily associated with chronic inflammation and fibrosis.[9,24–27] Inflammation is primarily aggravated by an increase in proinflammatory cytokine release, reactive oxygen species, and production of surface adhesion molecules.[26] These events lead to the migration of immune cells through renal tissue and activate fibrotic factors which subsequently cause fibrosis.[2] Our group and others shows that continuous activation of NKA signaling function leads to pathological effects like chronic inflammation and fibrosis.[2,19,26,28,29] Recent studies have demonstrated that 20-HETE may also interact with NKA, and is associated with development of CKD.[5,30,31] As a result, we aimed to investigate the specific molecular mechanisms of 20-HETE and NKA interaction through *in silico* analysis. Our objective was to better understand whether 20-HETE was capable of direct interaction with the NKA comparable to known NKA ligands such as CTS.

Re-docking is a pivotal validation step in the assessment of molecular docking methodologies. In our study, we initially attempted to dock Ouabain by directly extracting it from the PDB structure and docking it with its corresponding NKA structure. However, the obtained superimposed conformer RMSD value was 2.5 Å (**Figure S2**). The main reason is Ouabain has a large sugar group at the C-3 position of its steroidal structure, and forms a significant number of rotatable bonds, resulting in a high degree of molecular flexibility. Therefore, we used Ouabagenin (CTS without the sugar moiety) to dock with its own NKA crystal structure. This led to a superimposed ligand RMSD value of 0.85 Å, reflecting a more reliable and consistent representation of the ligand binding conformation within the active site of NKA.

To contextualize docking scores, a chemically diverse previously published dataset was screened to establish a background distribution of binding affinities toward Na^+^/K^+^-ATPase. This distribution reflects the range of non-specific interactions expected across general chemical space. Mapping biologically relevant ligands onto this distribution revealed that CTS, a known high-affinity ligands, occupy the high affinity region, whereas the negative controls DMSO and EtOH, included because they are commonly used solvents for dissolving 20-HETE and CTS in experimental studies, localize to the weak-binding region of the pocket. Notably, 20-HETE resides within an intermediate region of the distribution, supporting its potential to interact with NKA while maintaining consistency with its known biological role.

When a ligand binds to a protein, it triggers significant conformational changes. These changes are crucial to consider during molecular docking analysis since the ligand and protein adapt to each other as they form a complex.[32] Accurately predicting these protein conformational changes upon ligand binding remains a significant challenge. To address this, Schrödinger developed the IFD method, which mimics the induced fit mode of protein-ligand docking.[33] IFD experiments were conducted, introducing flexibility to the protein side chains, allowing the ligand to fine-tune its binding interactions within the active site, thus ensuring a better fit between the active site and the ligand.

Our rigid docking and IFD analyses consistently demonstrated a similar trend in binding affinity, showing that 20-HETE exhibits greater predicted affinity for NKA than negative controls, although lower than CTS. Notably, a difference between rigid docking and IFD were observed in this study. In the rigid docking analysis, the binding affinity order is as follows: MBG=110.2, TCB= −10.1, Ouabain=10.0, and Ouabagenin= −8.5. However, in the induced fit docking, Ouabain exhibits greater binding affinity with a score of −13.2, followed by Ouabagenin > TCB > MBG in the score respectively −10.6 > −10.2 > −10. These differences can be explained by the increased reliability associated with IFD, where it excels in capturing minor changes in molecular conformations. Consequently, IFD offers a more comprehensive and robust approach for ligand binding prediction, allowing for a finer exploration of conformational flexibility compared to rigid docking.

Further, in IFD, Ala330 was predicted to form a non-classical methyl C–H···O interaction with the hydroxyl oxygen of 20-HETE, with an H···O distance of 2.04 Å and a C–H···O bond angle of 146.9°, consistent with previously reported methyl C-O interactions in protein structures **(Figure S3)**.

Our analysis of binding free energy data reveals essential insights into the significance of specific amino acids in the interaction between 20-HETE and NKA. Notably, the mutated complexes generally exhibit lower binding free energies compared to the native NKA complex. However, Ala330Gly (−20.2651 Kcal/mol) and Val329Ala (−20.6945 Kcal/mol) have reduced the binding free energy to less than half a Kcal/mol than their native counterpart, and Thr804Ala has a binding free energy of −30.466 Kcal/mol, which is 13.4896 Kcal/mol less than the native complex. This underscores the importance of these amino acids in facilitating interaction with ligands. However, it is interesting to note that the Ile327 mutation yields a binding free energy of 42.203 Kcal/mol, which is similar to the native’s −43.9556 Kcal/mol. This phenomenon could be explained by the spatial distance between 20-HETE and Ile327, which might be greater than that between Ile328 and the ligand. Additionally, the hydrophobic interactions between these amino acids and 20-HETE imply that the strength of interaction diminishes with increased distance (**Figure S4**).

The mutation of Phe790 to Ala surprisingly resulted in an enhanced binding affinity from −43.9556 Kcal/mol to −46.7048 Kcal/mol. The bulky side chain of phenylalanine (Phe) is substantially larger compared to the alanine (Ala) side chain, which raised the possibility that bulkiness of phenylalanine side chain may have hindered interactions with 20-HETE. However, the mutation to Ala could have created space, potentially allowing 20-HETE to interact with other neighboring amino acids in close proximity **(Figure S5)**. This observation underscores the significance of Phe790 as an intriguing target for further investigation, especially regarding potential clinical implications. This mutation could cause persistent interactions between 20-HETE and NKA. Further structural and experimental studies will be required to understand the functional relevance of this observation. The results of our study provide critical insights into the molecular interactions between 20-HETE and the NKA, particularly emphasizing the relevance of specific amino acids in this interaction. Our binding affinity analysis, both through rigid docking and IFD, revealed that 20-HETE exhibits a notable affinity for NKA, similar to known CTS. The significance of specific amino acids was highlighted through mutational analysis, where mutations like Ala330Gly and Val329Ala drastically reduced binding free energy, indicating their critical role in 20-HETE binding. Additionally, the unexpected enhancement of binding affinity with the Phe790Ala mutation suggests that spatial constraints play a crucial role in ligand interaction, opening avenues for further exploration of the clinical implications of these findings, particularly in the context of CKD. This work underscores the potential for targeting specific NKA-ligand interactions in therapeutic strategies and provides a foundation for further investigation into the role of 20-HETE in CKD pathogenesis.

## 4. Materials and Methods

### 4.1 Protein structure selection and preparation for docking

The NKA structure with Protein data bank (PDB) ID 3A3Y was selected for this study. The NKA crystal structure is co-crystalized with Ouabain in E2P state with 2.8 Å resolution.[19] The structure was prepared using the protein preparation wizard in Maestro (Schrodinger,2022). The preprocessing procedure is followed as Allegra et al[17] and Pattar et al.[34] During preprocessing the Hydrogen atoms and bond order were assigned to the structure by OPLS2005 forcefield. The missing crystalized residues and missing atoms were fixed using the Schrodinger Prime™ tool, which is a fully integrated protein structure prediction solution that incorporates both homology modeling and fold recognition. Hydrogen bond assignment was optimized using PROPKA. Co-crystallized water molecules beyond 5.0 Å were removed and the structure was energy minimized.

### 4.2 Ligand preparation

A previously published dataset of chemically diverse compounds with no reported Na^+^/K^+^-ATPase binding was used to define a baseline distribution of docking scores.

To this investigation, CTS were chosen as positive control compounds, while 20-HETE and its analog were selected as experimental compounds of primary interest. Additionally, DMSO and EtOH were incorporated as negative control into the study. Ligand preparation involved Schrodinger Maestro LigPrep with OPLS2005 forcefield and EpiK ionization tool. Tautomer generation was allowed, and a maximum of 32 conformers was generated. Electrostatic potential values, determined by ESP atomic charges with OPLS2005, were mapped onto the ligand surfaces.

### 4.3 Structural validation and Rigid docking using Glide

The binding affinity of the molecules was assessed using the Maestro Glide tool by Schrodinger (2022). To validate the structural integrity post-preparation, a re-docking procedure was carried out, involving the extraction of the co-crystallized ligand from the PDB structure. This ligand was then docked into the NKA structure (3A3Y), and the resulting docked structure was superimposed with the crystal structure. Following this validation step, rigid docking was extended to include all the target ligands. The receptor grid was established with its center positioned at the co-crystallized ligand, utilizing a standard grid size. Throughout the ligand docking simulations, specific parameters were applied, including the scaling of van der Waals radii with a scaling factor of 0.8 and a partial charge cutoff set at 0.15. The XP (extra precision docking module) was used, with flexible ligand sampling, and no constraints were imposed during the docking process. After the docking simulations, a post-docking minimization process was carried out, considering ten poses per ligand, and a threshold of 0.5 Kcal/mol was applied to reject minimized poses that did not meet the specified criteria.

### 4.4 Induced fit docking

The Induced Fit Docking (IFD) procedure followed a previously established protocol.[17] In summary, all of our target ligands were employed in the IFD, with no constraints applied. The initial docking process used the co-crystallized ligand Ouabain as a centroid in the prepared structure. A van der Waals scaling factor of 0.5 was applied to both the receptor and ligand during this initial docking step, with a maximum of 20 ligand poses selected. In the subsequent Prime refinement step, side chains of residues within a 5 Å radius of the ligand were considered. In the redocking phase, XP precision docking was employed, with a maximum of 20 poses, and structures with an energy of 30 kcal/mol or less were redocked. For each ligand docked maximum 20 poses were retained to be redocked at the XP mode.

Subsequently, we examined the interaction fingerprint between 20-HETE and NKA via IFD simulations. We then identified specific amino acids of NKA as target candidates, mutated them, and conducted a comparative assessment of the binding free energy for each mutant.

### 4.5 Prime-molecular mechanics based generalized Born/surface area (MM-GBSA) calculation of free energy

Mutated receptors were generated using Maestro, followed by induced fit docking for each mutant and the ligand 20-HETE, as previously described.[17] The Schrodinger Prime module was employed to calculate the MM/GBSA relative binding free energy of the ligand-receptor complexes. For both native and mutated protein-ligands. The following equation was utilized for the calculation. [22]

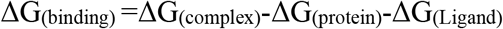

Here ΔG_(binding)_ is the binding free energy and ΔG_(complex)_, ΔG_(protein)_ and ΔG_(Ligand)_ are the binding free energy of complex, protein, and ligand respectively.[22]

In this study to calculate the relative binding free energy of native and each mutated protein complex, OPLS4 forcefield, default minimization steps were chosen and VSGB solution model was used as a solvent model.

The summary method pipeline is illustrated in **Figure 6**, detailing the overall process of molecular modeling work using Schrodinger’s small drug tool.

**Figure 6.**
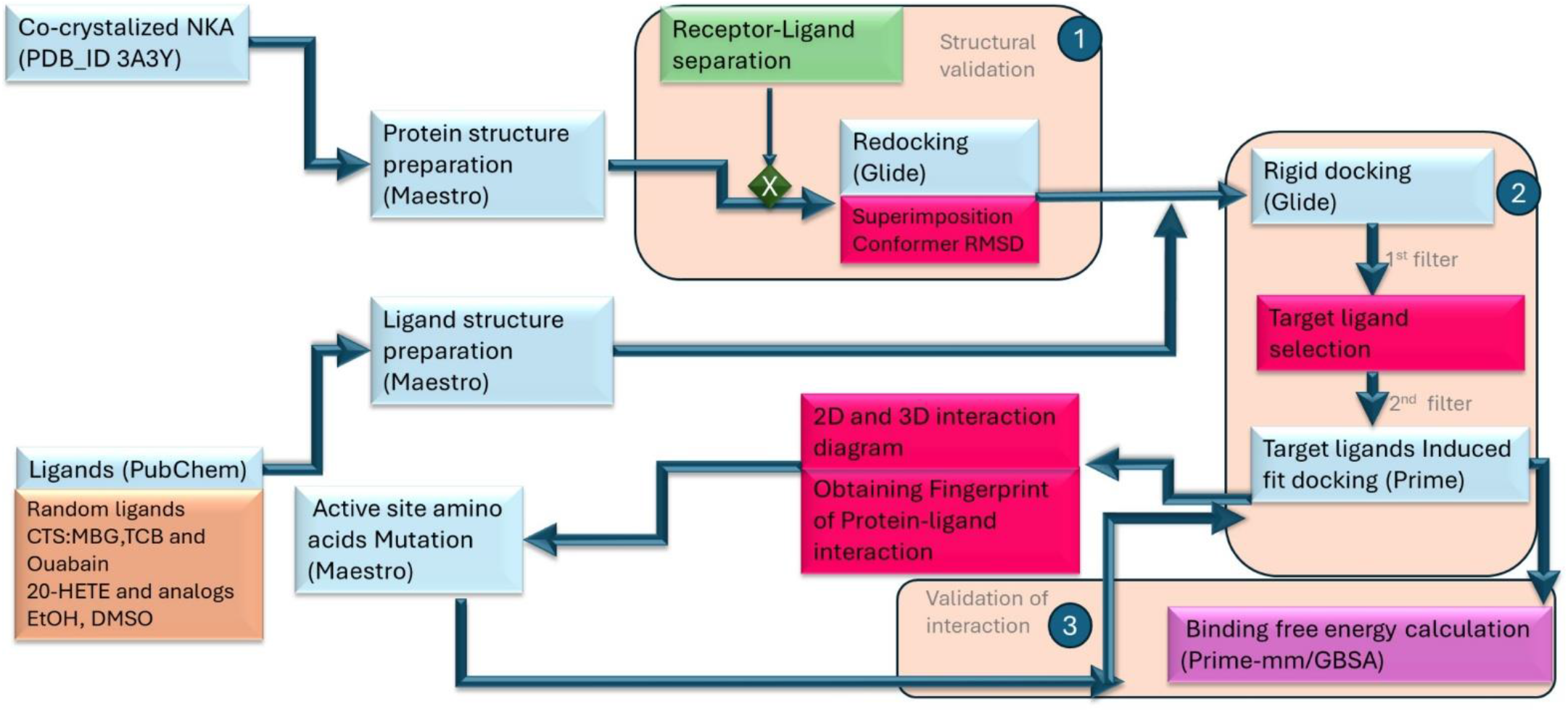
Method pipeline for the overall process of molecular modeling work using Schrodinger small drug tool

## 5. Conclusions

In conclusion, our *In-silico* study suggests that 20-HETE can potentially interact and bind with NKA, within the canonical ligand binding pocket. similar to CTS, although with less affinity. Rigid docking, IFD, interaction fingerprint analysis, and the binding energy calculations suggest that Ala330, Val329, and Thr804 as the key amino acids that may contribute to ligand stabilization. These findings collectively shed light on the dynamic interactions underlying 20-HETE binding to NKA and provide valuable insights into the potential binding mechanisms.

Nevertheless, additional biochemical and functional validation are required to confirm the findings from this theoretical study.

## Author Contributions

Conceptualization and Investigation, D.F., D.J.K.; methodology, D.F., R.A.; software, D.F, R.A, D.J.K.; validation, D.F; data curation and Formal analysis, D.F., R.A.,S.B.; resources, D.F.,R.A.; writing-original draft preparation, D.F., R.A.; writing-review and editing, D.F.,V.A.,S.T.H., D.J.K.; visualization, D.F, V.A.B.A.; supervision., D.J.K., S.T.H.; project administration, D.J.K, S.T.H.; funding acquisition, D.F., D.J.K., S.T.H. All authors have read and agreed to the current version of the manuscript.

## Funding

The authors disclose support for this work from the National Institutes of Health (HL-137004, D.J.K., S.T.H., the David and Helen Boone Foundation Research Fund (D.J.K.), the University of Toledo Women and Philanthropy Genetic Analysis Instrumentation Center (D.J.K., S.T.H.), a University of Toledo Graduate Student Association Award (D.F.), and the State of Ohio Supercomputer Center (D.J.K., D.F.)

## Institutional Review Board Statement

Not applicable

## Data Availability Statement

Data will be available upon request.

## Acknowledgments

The work was supported by Ohio Supercomputer Center for providing essential computational power.

## Conflicts of Interest

The authors declare no conflicts of interest.

## Supplementary Materials

**Figure S1.**
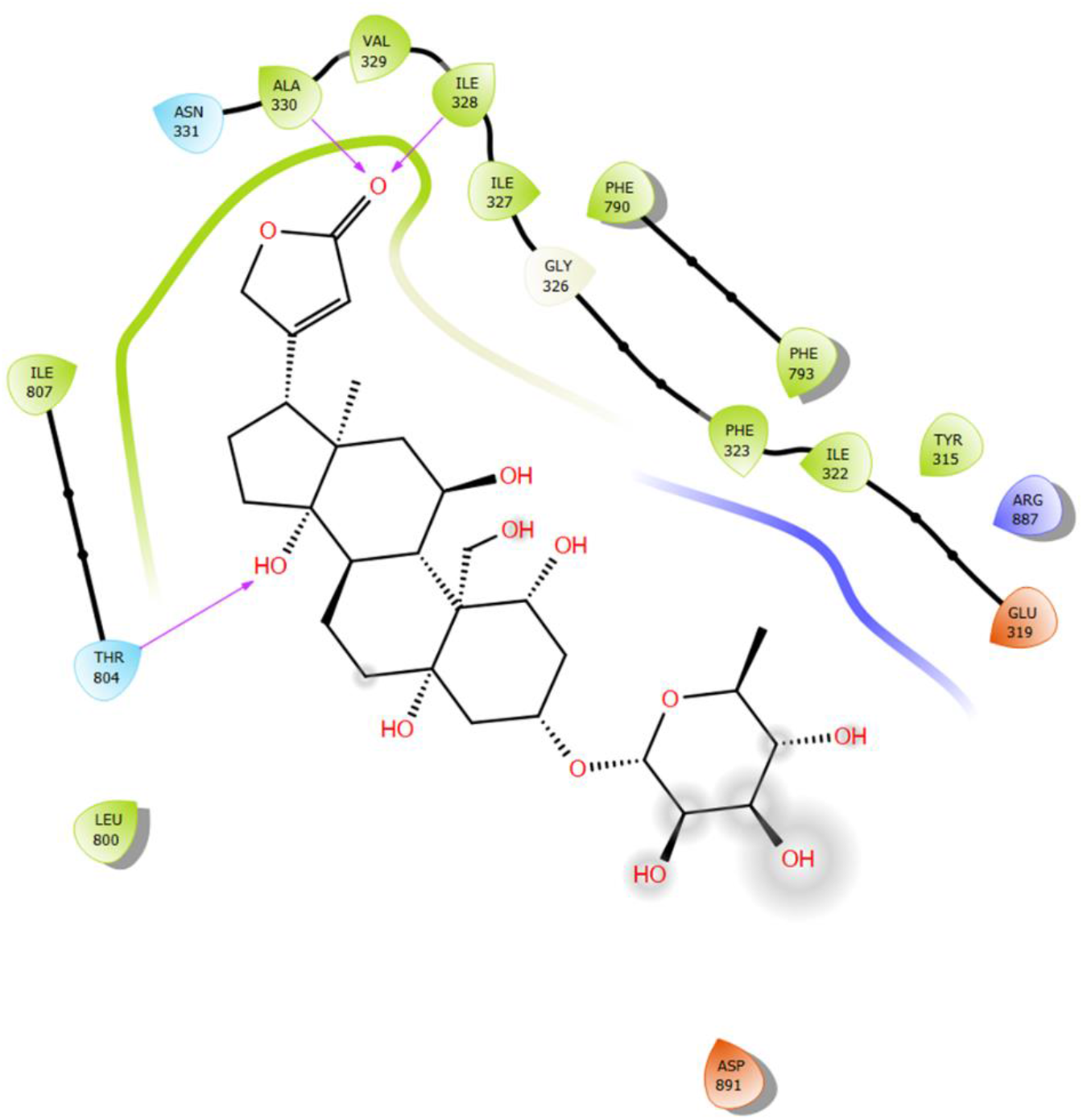
2D interaction diagram of the co-crystallized Ouabain within the binding pocket of Na+/K+-ATPase (3A3Y) generated using Schrödinger Maestro.

**Figure S2.**
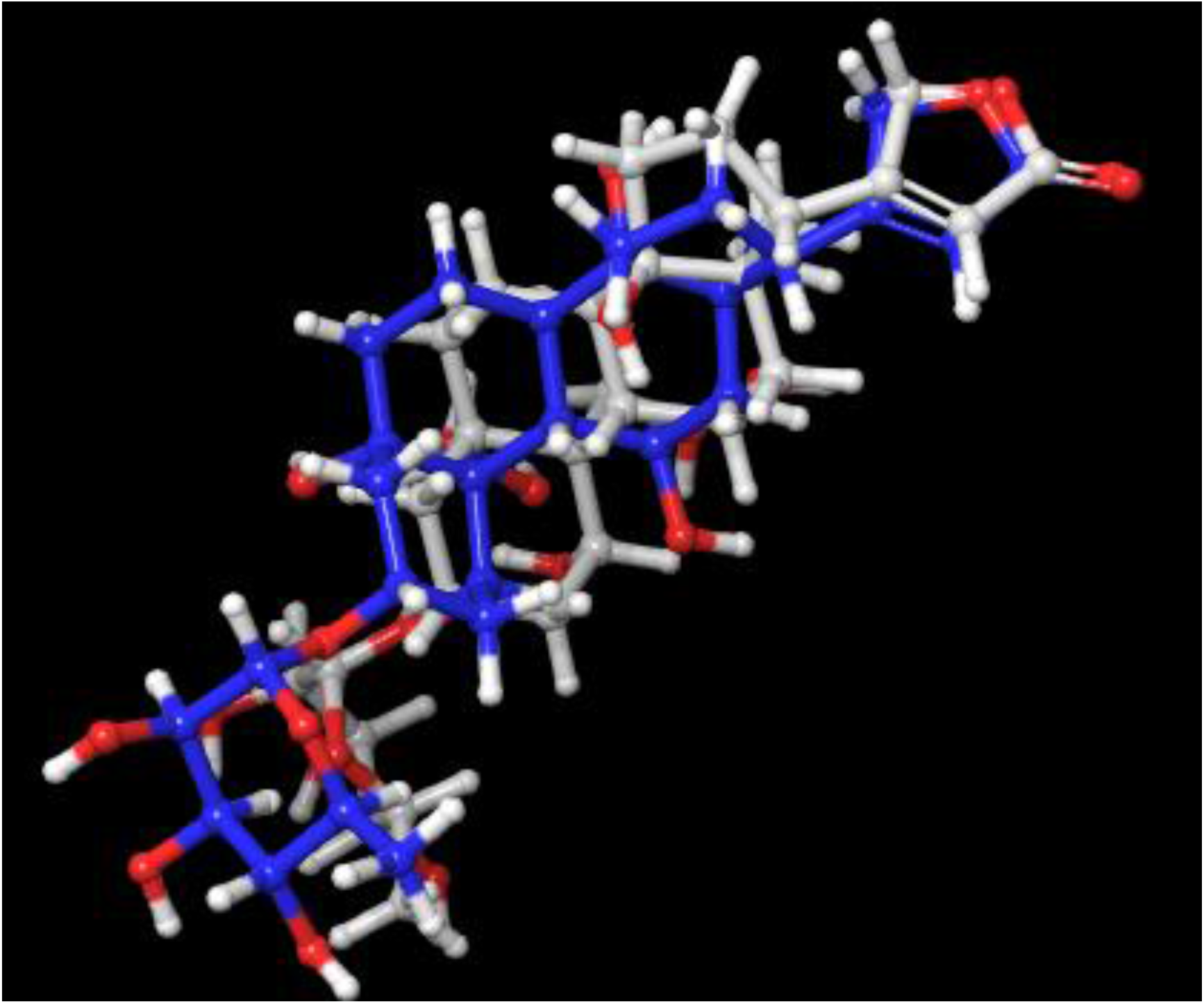
Superimposed 3D view of redocked Ouabain with its co-crystallized Ouabain. The co-crystallized ligand is shown in blue, and the docked ligand is shown in grey

**Figure S3.**
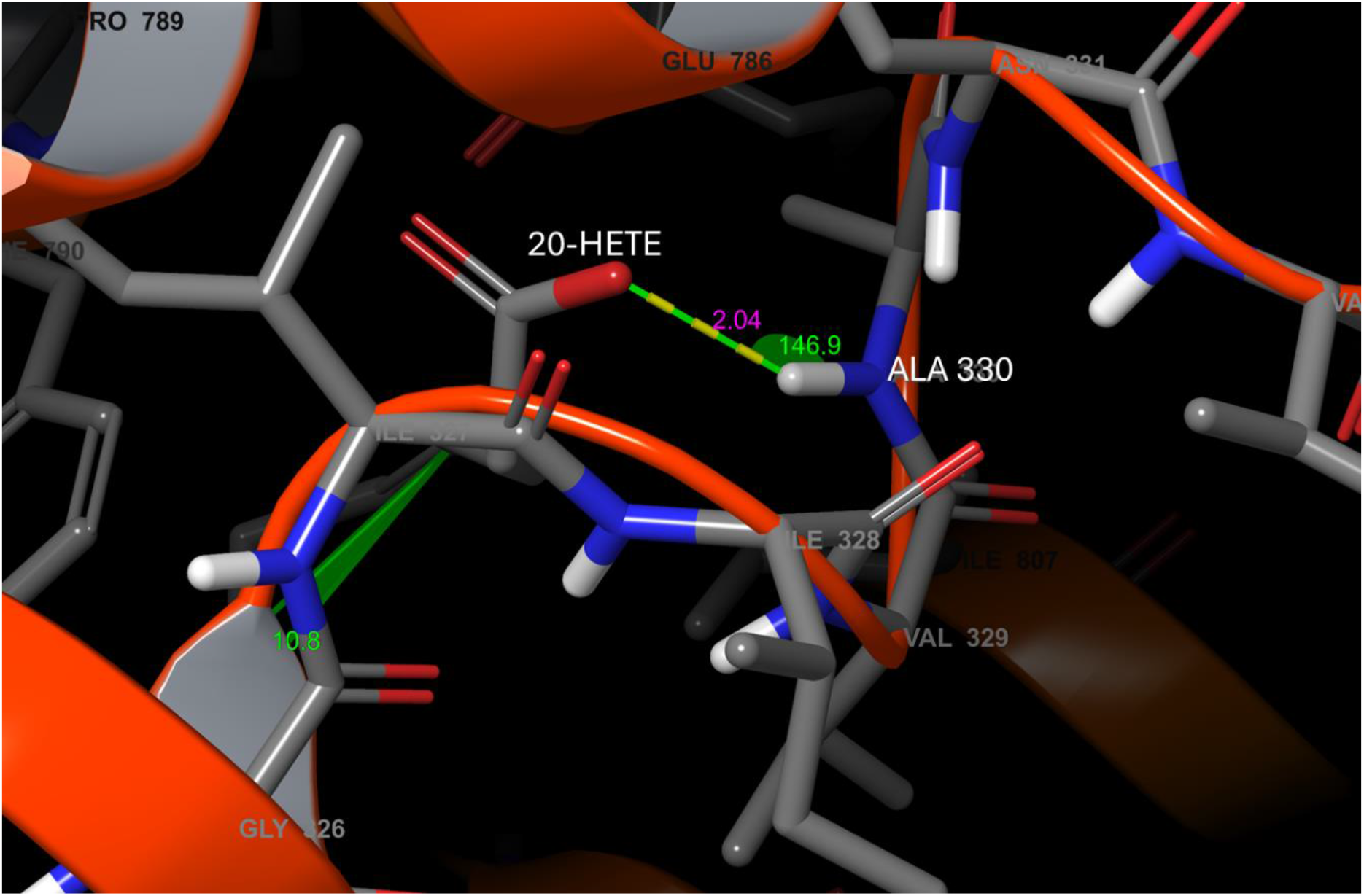
The predicted interaction geometry showed an H···O distance of 2.04 Å and a C–H···O bond angle of 146.9° between NKA, ALA 330 (blue and gray in color) and 20-HETE hydroxyl group (red in color)

**Figure S4.**
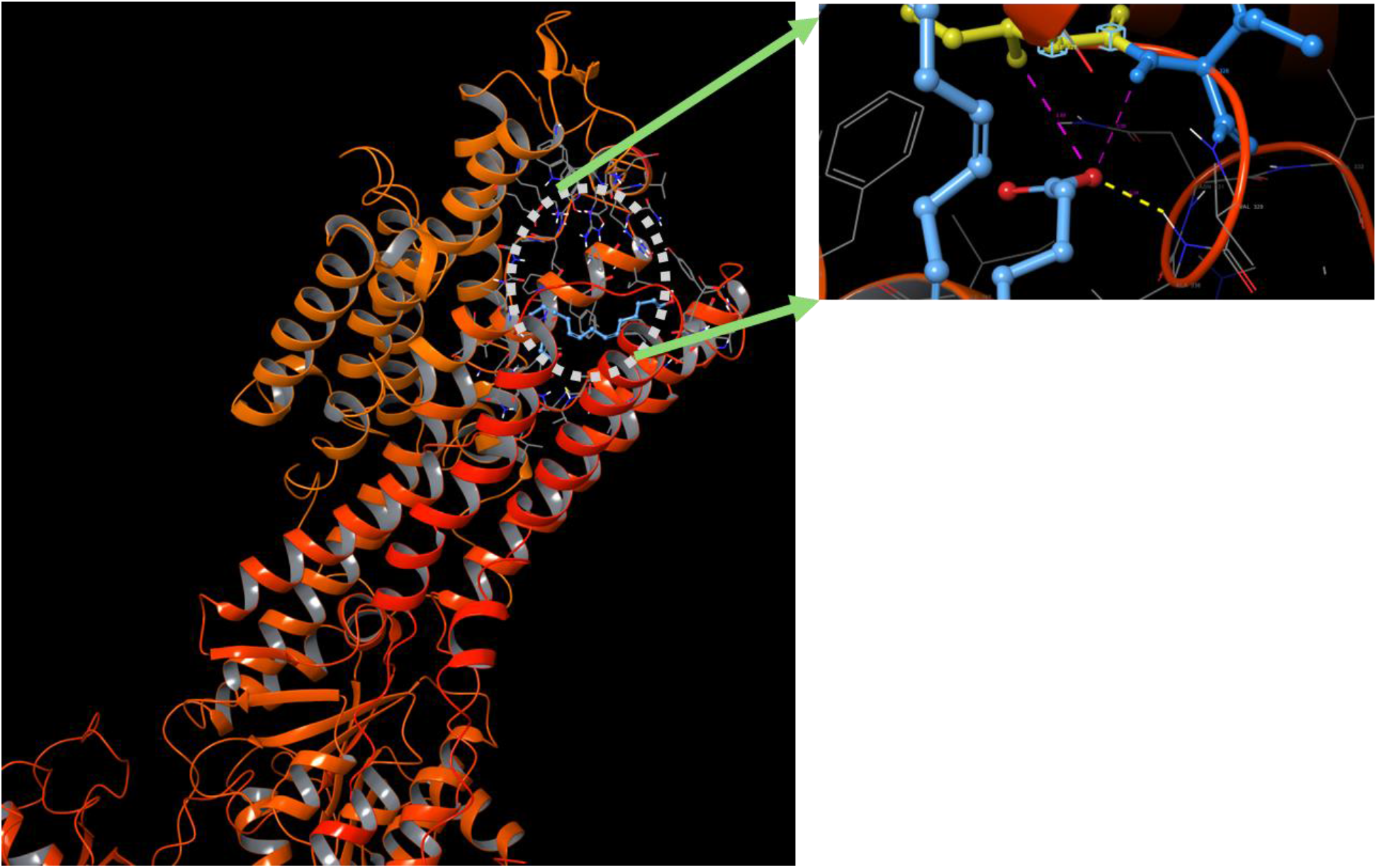
Spatial arrangement of Ile 327 and Ile 328, and the Distance to 20-HETE. Cyan representing 20-HETE and yellow indicating Ile 327 and light blue indicating Ile328 of NKA

**Figure S5.**
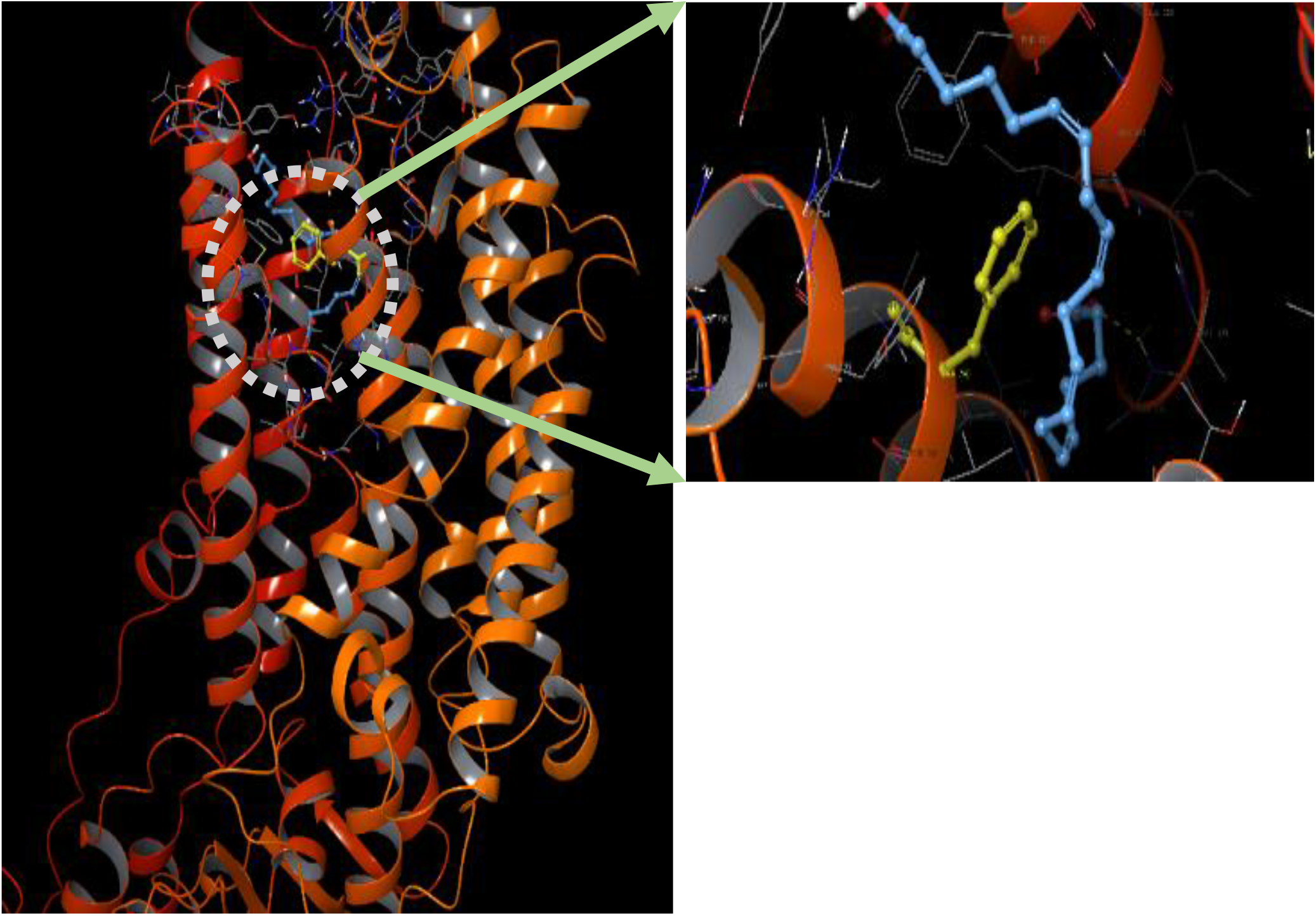
Spatial arrangement of Phe 790 and 20-HETE. Cyan representing 20-HETE and yellow indicating Phe 790

## Notes

### Competing Interest Statement

The authors have declared no competing interest.

## References

1. Cui, X.; Xie, Z. Protein Interaction and Na/K-ATPase-Mediated Signal Transduction. Molecules 2017, 22, doi:10.3390/molecules22060990.

2. Khalaf, F.K.; Dube, P.; Mohamed, A.; Tian, J.; Malhotra, D.; Haller, S.T.; Kennedy, D.J. Cardiotonic Steroids and the Sodium Trade Balance: New Insights into Trade-off Mechanisms Mediated by the Na+/K+-ATPase. Int. J. Mol. Sci. 2018, 19, 11–13, doi:10.3390/ijms19092576.

3. Rui, H.; Artigas, P.; Roux, B. The Selectivity of the Na+/K+-Pump Is Controlled by Binding Site Protonation and Self-Correcting Occlusion. Elife 2016, 5, 1–24, doi:10.7554/eLife.16616.

4. Al, X.S. et HHS Public Access. Physiol. Behav. 2017, 176, 139–148, doi:10.1002/iub.1229.Regulation.

5. Pratt, R.D.; Brickman, C.R.; Cottrill, C.L.; Shapiro, J.I.; Liu, J. The Na/K-ATPase Signaling: From Specific Ligands to General Reactive Oxygen Species. Int. J. Mol. Sci. 2018, 19, doi:10.3390/ijms19092600.

6. Bagrov, A.Y.; Shapiro, J.I.; Fedorova, O. V. Endogenous Cardiotonic Steroids: Physiology, Pharmacology, and Novel Therapeutic Targets. Pharmacol. Rev. 2009, 61, 9–38, doi:10.1124/pr.108.000711.

7. Kennedy, D.J.; Khalaf, F.K.; Sheehy, B.; Weber, M.E.; Agatisa-Boyle, B.; Conic, J.; Hauser, K.; Medert, C.M.; Westfall, K.; Bucur, P.; et al. Telocinobufagin, a Novel Cardiotonic Steroid, Promotes Renal Fibrosis via Na+/K+-Atpase Profibrotic Signaling Pathways. Int. J. Mol. Sci. 2018, 19, doi:10.3390/ijms19092566.

8. Ominato, M.; Satoh, T.; Katz, A.I. Regulation of Na-K-ATPase Activity in the Proximal Tubule: Role of the Protein Kinase C Pathway and of Eicosanoids. J. Membr. Biol. 1996, 152, 235–243, doi:10.1007/s002329900101.

9. Zhou, Y.; Yu, J.; Liu, J.; Cao, R.; Su, W.; Li, S.; Ye, S.; Zhu, C.; Zhang, X.; Xu, H.; et al. Induction of Cytochrome P450 4A14 Contributes to Angiotensin II-Induced Renal Fibrosis in Mice. Biochim. Biophys. Acta - Mol. Basis Dis. 2018, 1864, 860–870, doi:10.1016/j.bbadis.2017.12.028.

10. Hoopes, S.L.; Garcia, V.; Edin, M.L.; Schwartzman, M.L.; Zeldin, D.C. Vascular Actions of 20-HETE. Prostaglandins Other Lipid Mediat. 2015, 120, 9–16, doi:10.1016/j.prostaglandins.2015.03.002.

11. Ishizuka, T.; Cheng, J.; Singh, H.; Vitto, M.D.; Manthati, V.L.; Falck, J.R.; Laniado-Schwartzman, M. 20-Hydroxyeicosatetraenoic Acid Stimulates Nuclear Factor-KB Activation and the Production of Inflammatory Cytokines in Human Endothelial Cells. J. Pharmacol. Exp. Ther. 2008, 324, 103–110, doi:10.1124/jpet.107.130336.

12. Fan, F.; Roman, R.J. Effect of Cytochrome P450 Metabolites of Arachidonic Acid in Nephrology. J. Am. Soc. Nephrol. 2017, 28, 2845–2855, doi:10.1681/ASN.2017030252.

13. Stępniewska, J.; Dołęgowska, B.; Puchałowicz, K.; Gołembiewska, E.; Ciechanowski, K. Bioactive Lipids Derived from Arachidonic Acid Metabolism in Different Types of Renal Replacement Therapy. Chem. Phys. Lipids 2017, 206, 71–77, doi:10.1016/j.chemphyslip.2017.05.003.

14. Methods, D.; Design, L.; Data, V.; In, S.; Era, S.G. Docking Methods, Ligand Design, and Validating Data Sets in the Structural Genomics Era. 635–664.

15. Miller, E.B.; Murphy, R.B.; Sindhikara, D.; Borrelli, K.W.; Grisewood, M.J.; Ranalli, F.; Dixon, S.L.; Jerome, S.; Boyles, N.A.; Day, T.; et al. Reliable and Accurate Solution to the Induced Fit Docking Problem for Protein-Ligand Binding. J. Chem. Theory Comput. 2021, 17, 2630–2639, doi:10.1021/acs.jctc.1c00136.

16. Sahu, I.D.; McCarrick, R.M.; Lorigan, G.A. Use of Electron Paramagnetic Resonance to Solve Biochemical Problems. Biochemistry 2013, 52, 5967–5984, doi:10.1021/bi400834a.

17. Allegra, M.; Tutone, M.; Tesoriere, L.; Attanzio, A.; Culletta, G.; Almerico, A.M. Evaluation of the IKKβ Binding of Indicaxanthin by Induced-Fit Docking, Binding Pose Metadynamics, and Molecular Dynamics. Front. Pharmacol. 2021, 12, 1–13, doi:10.3389/fphar.2021.701568.

18. Li, H.; Leung, K.S.; Wong, M.H.; Ballester, P.J. Correcting the Impact of Docking Pose Generation Error on Binding Affinity Prediction. BMC Bioinformatics 2016, 17, doi:10.1186/s12859-016-1169-4.

19. Sandtner, W.; Egwolf, B.; Khalili-Araghi, F.; Sánchez-Rodríguez, J.E.; Roux, B.; Bezanilla, F.; Holmgren, M. Ouabain Binding Site in a Functioning Na+/K+ ATPase. J. Biol. Chem. 2011, 286, 38177–38183, doi:10.1074/jbc.M111.267682.

20. Side, F.; Methyl, C.; Bonding, C.H. Analysis of Neutron Structures. 2016, 83, 403–410, doi:10.1002/prot.24724.Frequent.

21. Dasmahapatra, U.; Kumar, C.K.; Das, S.; Subramanian, P.T.; Murali, P.; Isaac, A.E.; Ramanathan, K.; Balamurali, M.M.; Chanda, K. In-Silico Molecular Modelling, MM/GBSA Binding Free Energy and Molecular Dynamics Simulation Study of Novel Pyrido Fused Imidazo[4,5-c]Quinolines as Potential Anti-Tumor Agents. Front. Chem. 2022, 10, 1–17, doi:10.3389/fchem.2022.991369.

22. Muddagoni, N.; Bathula, R.; Dasari, M.; Potlapally, S.R. Homology Modeling, Virtual Screening, Prime-Mmgbsa, Autodock-Identification of Inhibitors of Fgr Protein. Biointerface Res. Appl. Chem. 2021, 11, 11088–11103, doi:10.33263/BRIAC114.1108811103.

23. Yang, H.C.; Zuo, Y.; Fogo, A.B. Models of Chronic Kidney Disease. Drug Discov. Today Dis. Model. 2010, 7, 13–19, doi:10.1016/j.ddmod.2010.08.002.

24. Charles, C.; Ferris, A.H. Chronic Kidney Disease. Prim. Care - Clin. Off. Pract. 2020, 1–17, doi:10.1016/j.pop.2020.08.001.

25. CDC CDC 2019 CKD Fact Sheet. Cdc 2019, 1, 1–6.

26. Khalaf, F.K.; Tassavvor, I.; Mohamed, A.; Chen, Y.; Malhotra, D.; Xie, Z.; Tian, J.; Haller, S.T.; Westfall, K.; Tang, W.H.W.; et al. Epithelial and Endothelial Adhesion of Immune Cells Is Enhanced by Cardiotonic Steroid Signaling Through Na+/K+-ATPase-α-1. J. Am. Heart Assoc. 2020, 9, doi:10.1161/JAHA.119.013933.

27. Mihai, S.; Codrici, E.; Popescu, I.D.; Enciu, A.M.; Albulescu, L.; Necula, L.G.; Mambet, C.; Anton, G.; Tanase, C. Inflammation-Related Mechanisms in Chronic Kidney Disease Prediction, Progression, and Outcome. J. Immunol. Res. 2018, 2018, doi:10.1155/2018/2180373.

28. Khalaf, F.K.; Dube, P.; Kleinhenz, A.L.; Malhotra, D.; Gohara, A.; Drummond, C.A.; Tian, J.; Haller, S.T.; Xie, Z.; Kennedy, D.J. Proinflammatory Effects of Cardiotonic Steroids Mediated by NKA α-1 (Na+/K+-ATPase α-1)/Src Complex in Renal Epithelial Cells and Immune Cells. Hypertension 2019, 74, 73–82, doi:10.1161/HYPERTENSIONAHA.118.12605.

29. Xie, J.X.; Zhang, S.; Cui, X.; Zhang, J.; Yu, H.; Khalaf, F.K.; Malhotra, D.; Kennedy, D.J.; Shapiro, J.I.; Tian, J.; et al. Na/K-ATPase/Src Complex Mediates Regulation of CD40 in Renal Parenchyma. Nephrol. Dial. Transplant. 2018, 33, 1138–1149, doi:10.1093/ndt/gfx334.

30. Akbulut, T.; Regner, K.R.; Roman, R.J.; Avner, E.D.; Falck, J.R.; Park, F. 20-HETE Activates the Raf/MEK/ERK Pathway in Renal Epithelial Cells through an EGFR-and c-Src-Dependent Mechanism. Am. J. Physiol. - Ren. Physiol. 2009, 297, 662–670, doi:10.1152/ajprenal.00146.2009.

31. Yu, M.; Lopez, B.; Dos Santos, E.A.; Falck, J.R.; Roman, R.J. Effects of 20-HETE on Na+ Transport and Na+-K +-ATPase Activity in the Thick Ascending Loop of Henle. Am. J. Physiol. - Regul. Integr. Comp. Physiol. 2007, 292, 2400–2405, doi:10.1152/ajpregu.00791.2006.

32. Christoph, A.S. Accounting for Induced-Fit Effects in Docking: What Is Possible and What Is Not. Curr. Top. Med. Chem. 2011, 11, 179–191.

33. Sherman, W.; Day, T.; Jacobson, M.P.; Friesner, R.A.; Farid, R. Novel Procedure for Modeling Ligand/Receptor Induced Fit Effects. J. Med. Chem. 2006, 49, 534–553, doi:10.1021/jm050540c.

34. Pattar, S.V.; Adhoni, S.A.; Kamanavalli, C.M.; Kumbar, S.S. In Silico Molecular Docking Studies and MM/GBSA Analysis of Coumarin-Carbonodithioate Hybrid Derivatives Divulge the Anticancer Potential against Breast Cancer. Beni-Suef Univ. J. Basic Appl. Sci. 2020, 9, doi:10.1186/s43088-020-00059-7.

